# Role of vaccinia virus growth factor in stimulating the mTORC1-CAD axis of the *de novo* pyrimidine pathway under different nutritional cues

**DOI:** 10.1101/2024.07.02.601567

**Authors:** Lara Dsouza, Anil Pant, Blake Pope, Zhilong Yang

## Abstract

Vaccinia virus (VACV), the prototype poxvirus, actively reprograms host cell metabolism upon infection. However, the nature and molecular mechanisms remain largely elusive. Given the diverse nutritional exposures of cells in different physiological contexts, it is essential to understand how VACV may alter various metabolic pathways in different nutritional conditions. In this study, we established the importance of *de novo* pyrimidine biosynthesis in VACV infection. We elucidated the significance of vaccinia growth factor (VGF), a viral early protein and a homolog of cellular epidermal growth factor, in enabling VACV to phosphorylate the key enzyme CAD of the *de novo* pyrimidine pathway at serine 1859, a site known to positively regulate CAD activity. While nutrient-poor conditions typically inhibit mTORC1 activation, VACV activates CAD via mTORC1-S6K1 signaling axis, in conditions where glutamine and asparagine are absent. However, unlike its cellular homolog, epidermal growth factor (EGF), VGF peptide alone in the absence of VACV infection has minimal ability to activate CAD, suggestive of the involvement of other viral factor(s) and differential functions to EGF acquired during poxvirus evolution. Our research provides a foundation for understanding the regulation of a significant metabolic pathway, namely, *de novo* pyrimidine synthesis during VACV infection, shedding new light on viral regulation under distinct nutritional environments. This study not only has the potential to contribute to the advancement of antiviral treatments but also improve the development of VACV as an oncolytic agent and vaccine vector.

**Importance:** Our research provides new insights into how VACV alters the mTORC1-CAD signaling axis under different nutritional cues. The identification of how VACV regulates a major enzyme, CAD, within the *de novo* pyrimidine synthesis pathway, establishes a molecular mechanism for determining how VACV reshapes this essential pathway, necessary for facilitating efficient VACV replication. We further emphasize that, despite nutrient-poor conditions, which typically inhibit mTORC1 activation, VACV can stimulate mTORC1. We identify its early growth factor, VGF, as an important factor for this stimulation of mTORC1 and its downstream effector CAD, revealing a new mechanism for how VACV sustains mTORC1-CAD axis activation under these nutrient deficient conditions. This work provides fresh insights into the molecular mechanisms of mTORC1-CAD regulation, which has the potential to be utilized to enhance VACV as an oncolytic tool, vaccine vector and aid in the development of antiviral drugs.

## Introduction

Viruses lack metabolism and rely on host cells to provide metabolites and energy for biosynthesis and replication^1–3^. Conceivably, nutritional conditions of host cells affect viral replication, and many viruses co-opt host metabolism to benefit viral replication. For example, vaccinia virus (VACV) demonstrates a preference for glutamine over glucose to achieve efficient replication^4^. This preference stems from the fact that asparagine, crucial for viral protein synthesis, is synthesized from glutamine, in addition to exogenous supply^5^. As we begin to understand how different nutrients affect viral replication, it is important to recognize that cells live under diverse nutritional conditions in physiological contexts, rather than in a uniformly nutrient-rich medium commonly used in laboratory cell culture systems^6^. Given the diverse range of nutritional environments of cells in various physiological contexts, a significant yet neglected aspect of virus-host interactions is the influence of various nutrients in the environment on viral alteration of host metabolic pathways and how a virus responds^7, 8^. We use VACV, the prototype poxvirus with a large double-stranded DNA genome, to understand the viral-host metabolism interactions under different nutritional cues.

Despite the eradication of smallpox, poxviruses remain a significant public health threat^9, 10^. This is exemplified by the current mpox (formerly known as monkeypox) outbreak, accentuating the ongoing challenges posed by poxviruses^10, 11^. However, many poxviruses have also made significant contributions to fighting other diseases and advancing biotechnology, such as in vaccine development and oncolytic virotherapy^12^. Employed as the vaccine for eradicating smallpox, VACV not only serves as the prototype member of the poxvirus family, but it is also the most extensively studied member of this family^13^. Because of its close genomic resemblance to other orthopoxviruses with many encoded proteins that are almost identical among othopoxviruses, VACV remains highly relevant for studying other highly pathogenic poxviruses (mpox, smallpox)^14^.

VACV encodes two copies of a gene encoding the early viral protein, vaccinia growth factor (VGF), which shares homology with cellular epidermal growth factor (EGF) and transforming growth factor-alpha (TGF-α)^15–17^. Among the 118 early VACV genes, VGF is among the most highly transcribed^18^. VACV secretes VGF, which competes with cellular epidermal growth factor (EGF) to bind to the EGF receptor (EGFR), thereby activating EGFR signaling and inducing proliferative effects in infected cells, consequently facilitating virus spread^19^.

An important, yet under studied aspect of poxvirus-host interactions, which limits the potential to develop new prevention and therapeutic strategies, is the modulation of key pathways that regulate host metabolism during poxvirus infection to produce the energy and metabolites necessary for viral replication^2^. We and others have previously reported that VACV actively alters host cell metabolism^20–23^. We demonstrated elegant mechanisms that identified the requirement of VGF to increase TCA cycle intermediates via STAT3 and reprograming of fatty acid metabolism upon VACV infection^21, 23^. Another important aspect of metabolism is the realm of nucleotide metabolism as nucleotides serve as the building block for DNA and RNA synthesis^24^. Cells can produce nucleotides via either the *de novo* synthesis pathway or salvage pathways^25^. In non-proliferative resting cells, that do not require high amounts of metabolites, nucleotides are sourced from salvage pathways. These pathways essentially involve the recycling of nucleotides obtained from DNA and RNA degradation or the uptake of nucleosides or nucleobases from the extra cellular space^26, 27^. However, during increased demand for nucleotides, the *de novo* synthesis pathway comes into play and synthesizes nucleotides from amino acids and glucose for efficient virus replication^26, 27^.

Recent studies suggest that some viruses enhance the synthesis of nucleotides to facilitate replication^28–30^. Earlier research has demonstrated that VACV replication was reduced when treated with N-phosphonacetyl-L-aspartate (PALA), an inhibitor of the *de novo* pyrimidine pathway^31^. CAD (carbamoyl-phosphate synthetase II, aspartate transcarbamoylase and dihydroorotase), a key enzyme of the *de novo* pyrimidine pathway is known to be activated by upstream growth factor MAPK signaling^25, 32–34^. Given the need for a continuous supply of nucleotides to sustain efficient VACV replication and with increasing evidence of the role of growth factor signaling in enhancing pyrimidine synthesis, we hypothesize that VGF, VACV’s EGF homolog, is involved in the regulation of CAD upon VACV infection.

In this study, we focus on understanding how VACV modulates CAD under different nutrient environments. We focus on the effect of three nutrients present *in vitro*: FBS, along with two crucial amino acids, glutamine and asparagine, to examine their influence on the regulation of CAD during VACV infection. We report the importance of the *de novo* pyrimidine pathway for efficient VACV replication and demonstrate the role of its viral early protein, VGF, in activating a major enzyme CAD of this pathway through mTORC1 (mechanistic target of rapamycin complex I) under different nutrition cues. Emerged as a central regulator of metabolism, mTORC1 incorporates growth factor signals and nutrient availability to regulate both anabolic and catabolic processes^35, 36^. Understanding the diverse downstream impacts of mTORC1, particularly in the context of VACV infection, remains limited, despite its potential to substantially affect metabolic and growth-related processes. Our results offer a mechanistic insight into how diverse nutritional environments serve as cues for VACV to utilize/regulate signaling pathways differently, facilitating its efficient replication. VACV has been developed for oncolytic therapy and given that cancer cells are prone to becoming vulnerable under nutrient stress conditions, our study on the role of VGF in stimulating the mTORC1-CAD axis under different nutritional stimuli will not only be instrumental in developing therapeutic strategies against poxviruses but also in enhancing their effectiveness as oncolytic tools^37, 38^.

## Materials and Methods

### Cell culture

Human Foreskin Fibroblasts (HFFs) were maintained in Dulbecco’s minimal essential medium (DMEM; Fisher Scientific) and BS-C-1 cells (ATCC CCL-26) were maintained in Eagle’s minimal essential medium (EMEM; Fisher Scientific). Both media were supplemented with 10% fetal bovine serum (FBS; Peak Serum), 2 mM glutamine (VWR), 100 U/ml of penicillin, and 100 μg/ml streptomycin (VWR).

Specialized DMEM media (1x DMEM - no glucose, no glutamine, no phenol red; Fisher Scientific; Cat #A1443001) were used in indicated experiments. For the protocol involving 2% FBS with either 2 mM glutamine or 2 mM asparagine, HFFs were infected with WT-VACV or vΔVGF upon reaching confluence. The infections were carried out using 1x DMEM supplemented with 2% dialyzed FBS (Gibco) and either 2 mM glutamine (Quality Biological) or 2 mM asparagine (Sigma-Aldrich) as specified for 8 h at the indicated MOI. For starvation with 0.1% FBS with either 2 mM glutamine or 2 mM asparagine, HFFs were initially starved using 1x DMEM media supplemented with 0.1% dialyzed FBS and either 2 mM glutamine or 2 mM asparagine as specified for a period of 48 h followed by infection with WT-VACV or vΔVGF for 8 h with fresh media containing the same supplements as those used for starvation. For starvation without glutamine or asparagine, HFFs were initially starved for approximately 36 hours in 1x DMEM supplemented with 0.1% dialyzed FBS and 2 mM asparagine. Sixteen hours prior to infection, this media was replaced with 1x DMEM supplemented with 0.1% dialyzed FBS and glucose only, without asparagine or glutamine. The total starvation period for the HFF cells was 48 h, followed by infection for 8 h with fresh media containing the same supplements as those used for the 16 h starvation. All cells were cultured in a humidified incubator set at 37°C with 5% CO_2_.

### Viruses and infection

The VACV Western Reserve (WR) strain (ATCC VR-1354) and a mutant VACV with VGF gene deleted (vΔVGF) was generated previously^21^. The stage-specific recombinant VACVs, which express *Gaussia* luciferase under VACV early, intermediate, or late promoters (vEGluc, vIGluc, and vLGluc, respectively) were used in this study. Amplification, purification, and titration of these viruses were conducted following established protocols described elsewhere^39^. Virus infection was carried out using the indicated multiplicity of infection (MOI) in regular DMEM media supplemented with 2.5% FBS, 2 mM glutamine, and 1 g/L glucose (unless specified otherwise) followed by plaque assay.

### Plaque assay

BS-C-1 cells were infected with WT-VACV, and the media was replaced with EMEM supplemented with 0.5%methylcellulose (Fisher Scientific) 1 hpi. After 48 h, the media was discarded and replaced with 0.1% (w/v) crystal violet solution dissolved in 20% ethanol for 10 min. Plaques were counted and titers were determined using a previously described method^40^.

### Cell viability assay

The cell viability assay was conducted using the trypan blue exclusion method, following previously established procedures^41^. HFFs were cultured in a 12-well plate followed by trypsinization with 200 µl of trypsin, and consequently suspended in 500 µl of DMEM media. The cells were centrifuged at 1000 xg for 5 min at 4°C. The supernatant was discarded, and the pellet was resuspended in 30 µl of DMEM media. 10 µl of the suspended pellet was mixed with 10 µl of 0.4% trypan blue (VWR), and 10 µl of this mixture was added to the dual chamber counting slides and cell viability was measured using an automatic cell counter (Bio-Rad).

### *Gaussia* luciferase activity assay

A luminometer and the Pierce *Gaussia* luciferase flash assay kit (Thermo Scientific) were used to measure *Gaussia* luciferase activity per manufacturer’s instructions.

### Quantitative PCR

DNA was extracted utilizing the EZNA Blood DNA Kit. Relative levels of viral DNA were quantified through the CFX96 real-time PCR instrument (Bio-Rad, Hercules, CA) and the All-in-one 2x qPCR mix (GeneCopoeia), utilizing specific VACV primers targeting the C11R gene. As internal references, primers for the 18S rRNA gene were used. The initial denaturation step was started at 95.0°C for 10 min followed by 40 cycles of denaturation at 95°C for 10 s, annealing at 53°C for 30 s and extension at 72°C for 15 s. The sequence for the primers used are as follows:

C11p FW: AAACACACACTGAGAAACAGCATAAA

C11p Rev: ACTATCGGCGAATGATCTGATTATC

18S rRNA FW: CGA TGC TCT TAG CTG AGT GT

18S rRNA Rev: GGT CCA AGA ATT TCA CCT CT

### Inhibitors and antibodies

The chemical inhibitors leflunomide (Cat #1247), and rapamycin (Cat #s1039) were purchased from Selleck Chemicals. Pyrazofurin (Cat #SML1502) was purchased from Sigma Aldrich. PALA was acquired from the Drug Synthesis and Chemistry Branch, Development Therapeutics Program, Division of Cancer Treatment and Diagnosis, National Cancer Institute. Antibodies targeting CAD at serine1859 (Cat #12662), total CAD (Cat #11933), DHODH (Cat #80981), S6K1 T389 (Cat #9234), and total S6K1 (Cat #2708) were purchased from Cell Signaling Technology. CAD threonine 456 /pCPS2 (Cat #sc377559) and the anti-glyceraldehyde-3-phosphate dehydrogenase (anti-GAPDH) (Cat #sc-365062) antibody was sourced from Santa Cruz Biotechnology.

### Western blotting analysis

Western blotting analysis was conducted according to procedures described elsewhere^21^. Briefly, samples were prepared using NP-40 lysis buffer, followed by the addition of 100 mM dithiothreitol (DTT) as a reducing agent, and sodium dodecyl sulfate-polyacrylamide as the loading buffer. Following boiling at 95°C for 5 min, samples were loaded onto SDS–PAGE gels. The gels were subsequently transferred for 10 min onto a polyvinylidene difluoride membrane (PVDF) using the Trans-Blot turbo transfer system (Bio-rad). The membranes were blocked using blocking buffer prepared using 5% milk (Alkali Scientific) in 1x TBST for 1 h at room temperature, followed by overnight incubation with the primary antibody in blocking buffer (5% milk in 1x TBST) at 4°C or 1 h at room temperature. The membrane was washed three times for 10 min each with 1x TBST and incubated with a horseradish peroxidase-conjugated secondary antibody (Cell Signaling Technology; Cat #7074S) for 1 h at room temperature. Membranes were developed using Thermo Scientific SuperSignal West Femto Maximum Sensitivity Substrate (Thermo Fisher; Cat #34095) and imaged using Chemiluminescent western blot imager (Azure 300). Antibodies were stripped from the membrane using Restore Plus western blot stripping buffer (Fisher Scientific, Cat #46430) for Western blot analysis with another primary antibody.

### VGF peptide expression, purification, and treatment

The sequence for the cleaved, secreted portion of the VGF peptide was obtained from NCBI YP_232891.1^15–17^ A signal peptide was added at the N terminal region (underlined and was cleaved after secretion), and a 6xHis tag at the C terminal (<u>MGWSCIILFLVATATGVHS</u>DSGNAIETTSPEITNATTDIPAIRLCGPEGDGYCLHGDCIHARDIDGMYCRCSHGYTGIRCQHVVLVDYQRSENPNTHHHHHH) was expressed and purified using a mammalian expression system using the HD CHO-S cell line at GenScript. This process involved cloning the peptide’s secreted sequence with 6xHis tag encoded gene into the vector pcDNA3.4, followed by plasmid preparation, TurboCHO 2.0 expression, and one-step purification using Ni-NTA. The purity of the peptide was assessed by SDS-PAGE Coomassie Blue staining and SEC-HPLC. The peptide is stored in 1x PBS (pH 7.2).

### Statistical Analysis

The data presented represent the mean of at least three biological replicates, unless mentioned otherwise. Error bars depict the standard deviation of the experimental replicates. Students *t-test* was conducted to assess any significant differences between the two means. We utilized the following symbols to denote statistical significance: P > 0.05: ns, P ≤ 0.05: *, P ≤ 0.01: **, P≤ 0.001 ***. Figures were prepared using GraphPad version 10.2.3, and schematics were created using BioRender.com

## Results

### *De novo* pyrimidine synthesis is required for efficient VACV replication

The *de novo* synthesis of pyrimidines is initiated by the amino acid glutamine and the trifunctional enzyme CAD. The synthesis starts with the production of dihydroorotate followed by the oxidation of dihydroorotate to orotate by another enzyme, dihydroorotate dehydrogenase (DHODH). Orotate is further phosphorylated to produce uridine monophosphate (UMP) by bifunctional enzyme UMP synthase (UMPS). UMP is then utilized as the first metabolite for pyrimidine synthesis^42^**(Fig 1A).** Previous studies indicated that treatment with PALA, a chemical compound that selectively targets CAD, resulted in a substantial reduction in VACV replication in cultured cells^31, 43^. To further determine the significance of the *de novo* pyrimidine pathway during VACV infection, we used compounds selectively targeting the other two enzymes, DHODH and UMPS, of this pathway to examine their effects on VACV replication in HFFs^44–47^ **(Fig. 1A)**. In HFFs infected with VACV at an MOI of 2, we treated the cells with either leflunomide (a selective inhibitor of DHODH) or pyrazofurin (a selective inhibitor of UMPS) for 24 h. We observed a substantial and significant decrease in VACV titers compared to vehicle-treated samples, with reductions of 110-fold for cells treated with leflunomide and 16-fold for cells treated with pyrazofurin, respectively **(Fig 1B & 1D)**. These concentrations did not adversely affect the cell viability **(Fig 1C & 1E).** These findings further corroborate the importance of the *de novo* pyrimidine pathway during VACV replication.

**Figure 1.**
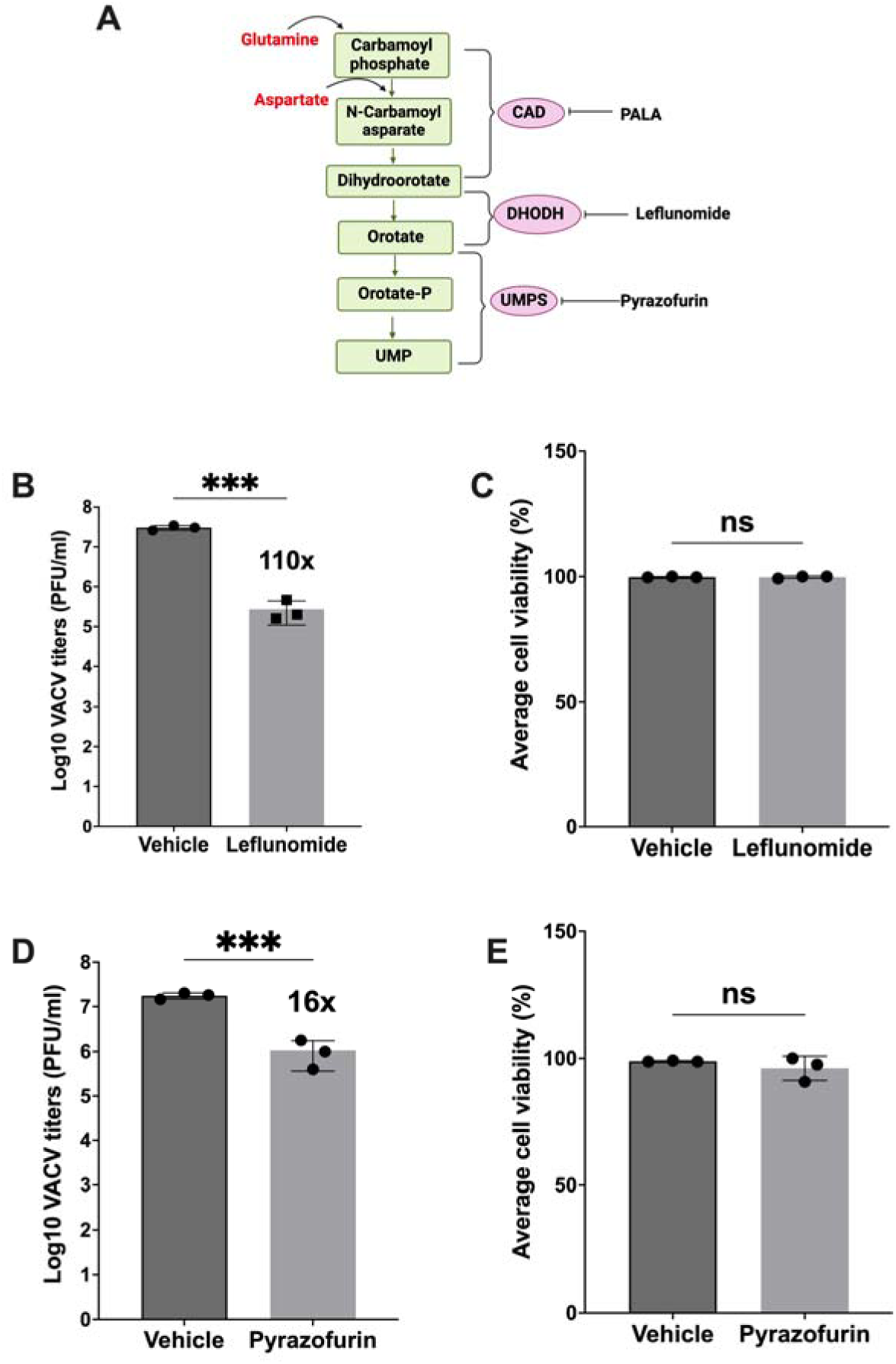
Inhibition of *de novo* pyrimidine pathway significantly reduces VACV replication. **(A)** An overview of the *de novo* pyrimidine pathway with compounds inhibiting key enzymes of this pathway. (**CAD** – carbamoyl phosphate synthetase II, aspartate transcarbamoylase, dihydroorotase) (**DHODH**-dihydroorotate dehydrogenase) (UMPS-uridine monophosphate synthetase) (**PALA** – N-phoshponacetyl-L-aspartate). **(B & D)** HFFs were infected with VACV at an MOI of 2 and treated with the indicated compounds 1 hpi at a concentration of 100 µM. Cell lysates were collected at 24 hpi for plaque assay using BSC-1 cells. **(C & E)** HFFs were treated with leflunomide or pyrazofurin (100 µM) and cell viability was measured using trypan blue staining after 48 h treatment. Error bars represent the standard deviation of at least three biological replicates. P> 0.05: ns, P ≤ 0.001:***. Statistical analysis for difference between two means were performed using the student’s T test.

### *De novo* pyrimidine pathway is required for VACV DNA replication and subsequent intermediate and late gene expression

VACV gene expression occurs in a temporal manner, dividing the expression of viral genes into three stages: early, intermediate, and late. After VACV entry, early genes are expressed^48^. Intermediate and late genes are expressed after DNA replication and requires the aid of transcription factors expressed from early and intermediate genes, respectively. To identify the stage of VACV replication affected by the inhibition of the *de novo* pyrimidine pathway, we used three recombinant VACVs with *Gaussia* reporter genes under the control of either an early promoter (C11R), intermediate promoter (G8R), or late (F17R) viral promoter **(Fig. 2A)**. Luciferase activities were measured at 8 hpi for intermediate and late gene expression and 4 hpi for early gene expression. While there was a subtle reduction in samples treated with leflunomide compared to the vehicle treated samples, we did not observe a substantial reduction in early gene expression between vehicle-treated and those treated with inhibitors in VACV-infected HFFs (MOI=2) **(Fig. 2B)**. A six-fold reduction was observed in intermediate gene expression of infected cells on treatment with inhibitors targeting enzymes CAD or DHODH **(Fig. 2C)**. A higher reduction in intermediate gene expression (∼1000-fold) was noted with the UMPS inhibitor **(Fig. 2C)**. Furthermore, a significant reduction (∼17-fold reduction for PALA, ∼26-fold reduction for leflunomide and ∼68-fold reduction for pyrazofurin) in late gene expression was observed upon inhibition of this pathway with each of the three inhibitors individually **(Fig. 2D)**. We additionally tested the effect of inhibiting the *de novo* pyrimidine pathway on VACV DNA amounts. Validating previous findings regarding the decrease in VACV DNA levels upon treatment with PALA, we observed a reduction of ∼3-fold in VACV DNA levels upon PALA treatment in HFF cells **(Fig. 2E)**. We also observed a ∼2-fold decrease in VACV DNA levels when treating infected HFFs with a DHODH inhibitor and a ∼7-fold reduction upon inhibition of UMPS **(Fig. 2F & 2G)**. As VACV intermediate and late gene expression relies on viral DNA synthesis, the reduction in intermediate and late gene expression could be partly because of lower viral DNA levels and/or slowed viral RNA synthesis. Nonetheless, the results indicate *de novo* pyrimidine pathway is required for VACV replication starting at the DNA replication stage.

**Figure 2.**
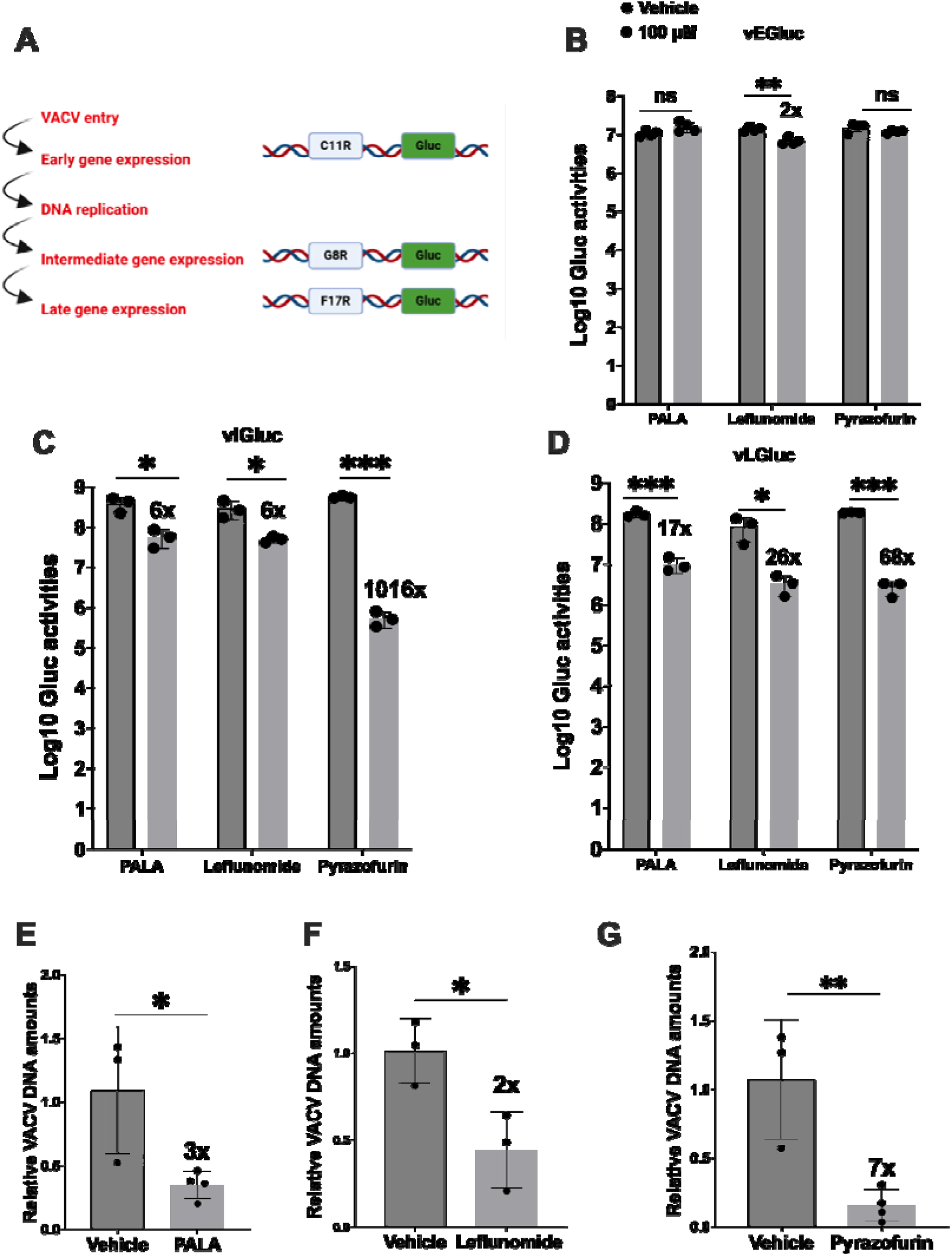
*De novo* pyrimidine pathway is required for VACV DNA replication and subsequent intermediate and late gene expression. **(A)** Schematic of the temporal expression pattern of VACV genes with recombinant VACV containing stage specific *Gaussia* reporter genes. **(B, C & D)** Inhibition of *de novo* pyrimidine pathway reduces VACV intermediate and late gene expression. VACV gene expression was quantified by measuring *Gaussia* luciferase activities using reporter VACVs expressing *Gaussia* luciferase under early (vEGluc**, B**), intermediate (vIGluc**, C**), and late (vLGluc, **D**) promoters, respectively. HFFs were pre-treated with the indicated compounds for approximately 16 h prior to infection and subsequently infected with the indicated VACV at an MOI of 2, followed by treatment with the indicated compounds 1 hpi. *Gaussia* luciferase activities were measured at 4 hpi for early gene expression and 8 hpi for intermediate and late gene expression. **(E, F & G)** Inhibition of *de novo* pyrimidine pathway reduces VACV DNA levels. HFFs were pretreated with indicated pyrimidine synthesis inhibitors 16 h prior to infection and subsequently infected with WT-VACV virus at an MOI of 2, followed by treatment with indicated *de novo* pyrimidine pathway inhibitors after 1 hpi. Cell lysates were collected at 8 hpi. DNA was extracted and relative viral DNA levels were determined using real time PCR with VACV specific primers. Error bars represent the standard deviation of at least three biological replicates. P> 0.05: ns, P≤ 0.05: *, P ≤ 0.01: **, P ≤ 0.001:***. Statistical analysis between two means was performed using the student’s T test.

### VGF is necessary for the phosphorylation of CAD at serine1859 in glutamine and asparagine deprived media

Human CAD is a large multifunctional key enzyme with approximately 2,225 amino acids, catalyzing the three initial rate-limiting steps of the pyrimidine pathway^25^. CAD has four domains namely: glutamine amidotransferase (GATase), carbamoyl phosphate synthetase II(CPSIIase), dihydroorotase (DHOase), and aspartate transcarbamoylase (ATCase)^25, 42^**(Fig. 3A).** Multiple amino acids of this protein can be phosphorylated by different signaling, leading to distinct mechanisms of regulation for this large enzyme^24, 49^. CAD undergoes post-translational modifications at numerous sites, with phosphorylation at serine 1859 (located between the DHOase and ATCase domain) and threonine 456 (located in the CPSIIase domain) being well studied and known to stimulate CAD activity^34, 50, 51^ **(Fig. 3A)**. To examine how VACV infection regulates the *de novo* pyrimidine pathway, we examined protein phosphorylation levels of the key enzyme CAD responding to VACV infection. Phosphorylation of CAD at threonine 456 (T456) is facilitated by the MAPK signaling pathway upon activation of its upstream receptor EGFR, whereas phosphorylation at serine1859 (S1859) occurs through the activation of its upstream protein S6K1, which in turn is activated through the mTORC1 signaling pathway^42^. Given that VGF stimulates growth factor signaling, we examined if VGF plays a role in modulating CAD phosphorylation. To determine the necessity of VGF in activating CAD upon infection, we assessed the phosphorylation levels of these two phosphorylation sites with well-established functions, under two different cell culture media (asparagine + glucose or glutamine + glucose) in the presence of 2% FBS. Glutamine serves as a major carbon source in cellular metabolism, the depletion of which decreases VACV replication^4^. Asparagine, an amino acid whose synthesis exclusively depends on glutamine, was shown to be important for VACV protein synthesis, thereby rescuing the effects of glutamine deprivation on VACV replication^5^. Asparagine-replete media, independent of glutamine, therefore, serves as a system to study the effects of VACV on metabolism, eliminating the masking effects of glutamine. We first tested the role of VACV infection on CAD phosphorylation in two media: the commonly used medium during VACV infection (glucose + glutamine) and a medium that eliminates the masking effect of glutamine to help us examine the role of individual viral factors upon VACV infection (glucose + asparagine) *in-vitro*. To determine the role of VGF, we used a recombinant VACV with both copies of the VGF gene deleted (vΔVGF). We did not observe substantial changes in phosphorylation levels of CAD at T456 and S1859 in mock (uninfected), wildtype vaccinia (WT-VACV), and vΔVGF infections in the presence of 2% FBS and glutamine conditions. Though we observed a subtle increase in CAD S1859 phosphorylation levels for WT-VACV infection compared to mock in 2% FBS and asparagine conditions, we did not observe a substantial difference between WT-VACV and vΔVGF infections in this condition. **(Fig. 3B)**.

**Figure 3:**
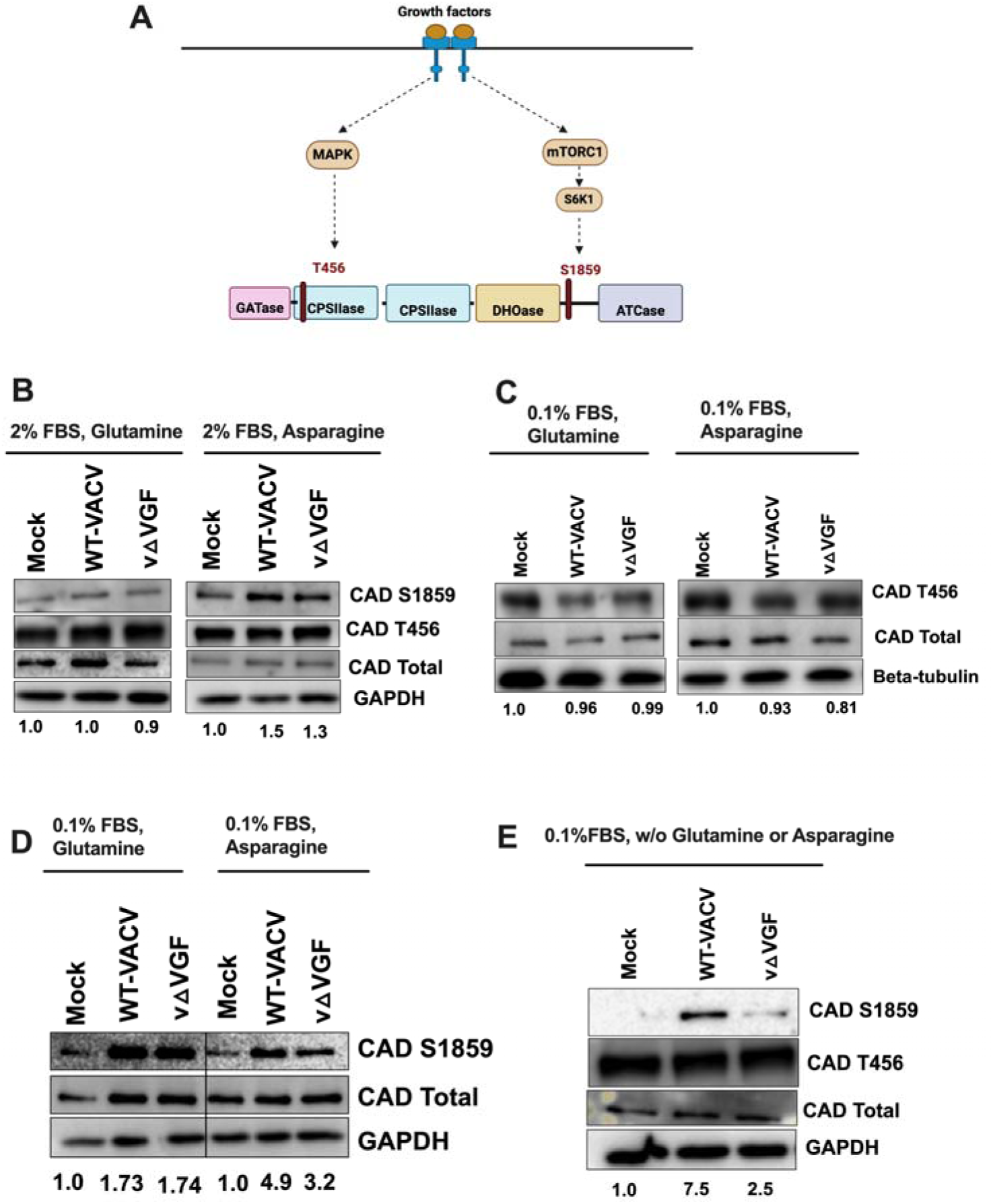
VGF activates CAD at S1859 in glutamine and asparagine depleted conditions. **(A)** Schematic representation of growth factor stimulation of CAD at two different phosphorylation sites. GATase: glutamine amidotransferase, CPSIIase: carbamoyl phosphate synthetase II, DHOase: dihydroorotase, ATCase: aspartate transcarbamoylase. **(B)** HFFs were mock-infected, infected with WT-VACV or v△VGF at an MOI of 5 with 2% dialyzed FBS, 2mM Glutamine or 2 mM Asparagine and cell lysates were collected for western blot sample preparation at 8 hpi. **(C&D)** HFFs were starved with 0.1% dialyzed FBS and 2 mM glutamine or 2 mM asparagine for 48 h prior to infection. Infected cell lysates were collected at 8 hpi. For (**C**), samples from both conditions were loaded on the same SDS-PAGE gel. **(E)** HFF’s were starved with 0.1% dialyzed FBS and 2mM asparagine, followed by replacement with media containing 0.1% dialyzed FBS lacking both glutamine and asparagine for approximately 16 h prior to infection. The infection media consisted of 0.1% dialyzed FBS without glutamine or asparagine. Infected cell lysates were collected at 8 hpi. Representative images from three biological replicates are shown. The numbers below the blots indicate the fold change of CAD S1859 (**B, D, & E**) and CAD T456 (**C**) based on the average intensities of Western blot analysis from at least three biological replicates, with each replicate normalized to the loading control of its respective blot, and then further normalized to its mock treatment. Western blot quantification was determined by NIH Image J.

To minimize the influence of factors that may be present in the FBS of cell culture media, we starved the HFF cells for a period of 48 h in the presence of minimal amount of FBS (0.1% dialyzed FBS) and either asparagine or glutamine. Upon starvation, we continued to observe similar levels of T456 phosphorylation in mock and infected (WT-VACV and vΔVGF) conditions in the presence of glutamine or asparagine **(Fig. 3C)**, while viral infection increased S1859 phosphorylation in the presence of either, glutamine or asparagine **(Fig. 3D)**. These data indicate that FBS, asparagine, and glutamine can individually contribute to CAD phosphorylation in the presence of VACV infection conditions, which may partially mask the effect of individual viral factors upon VACV infection. We observed a moderate, yet consistent decrease of CAD S1859 phosphorylation in vΔVGF infection compared to WT-VACV infection, in asparagine media with 0.1% FBS **(Fig. 3D)**, suggesting that VGF may stimulate CAD S1859 phosphorylation in nutrition-poor conditions.

To further determine the role of VGF in CAD S1859 activation upon viral infection, we examined it in media without glutamine and asparagine in the presence of minimal FBS (0.1%). We observed a greater reduction of CAD S1859 phosphorylation in starved HFFs in the absence of glutamine and asparagine, while VACV infection strongly elevated CAD S1859 phosphorylation by over 7.5-fold. There was a substantial decrease in the phosphorylation of CAD at S1859 in the vΔVGF infection condition compared to WT-VACV infection, indicating that glutamine or asparagine are capable of stimulating phosphorylation of CAD S1859 independent of VGF. There continued to be no obvious increase in the phosphorylation levels of CAD at T456 in conditions deprived of glutamine and asparagine indicative of VGF primarily playing a role in activating CAD through the S1859 site under nutrient poor conditions **(Fig 3E)**. These findings demonstrate differential responses of the key enzyme CAD to the viral protein VGF in the context of different nutrients *in vitro*, namely FBS, glutamine, and asparagine. This indicates the significance of VGF in promoting the *de novo* pyrimidine synthesis pathway in the context of nutritional deficiency, especially in conditions lacking glutamine and asparagine and/or growth signals. We did not observe changes in protein levels of DHODH, the second key enzyme of the *de novo* pyrimidine synthesis pathway, across various conditions (data not shown).

### VACV activates CAD S1859 through mTORC1-S6K1 in a VGF-dependent manner

We next sought to identify the signaling pathway responsible for phosphorylating CAD at S1859 during VACV infection. It has been previously identified that mTORC1, a master regulator of metabolism, could post-translationally modify CAD and increase its activity by phosphorylating it at S1859 through mTORC1’s downstream target, S6K1^50, 51^. To test if mTORC1 is important for activation of CAD in the context of VACV infection, we first examined if VGF is necessary for mTORC1 signaling by testing S6K1 phosphorylation (S6K1 T389), in different nutrient conditions^52^. We first determined the phosphorylation of S6K1 in the presence of glutamine or asparagine and minimal FBS (0.1%). With minimal FBS present in the media, there was a substantial increase in phosphorylation of S6K1 stimulated by VACV infection. However, there was no obvious difference between WT-VACV and vΔVGF conditions **(Fig. 4A & 4B)**. When depriving HFFs of glutamine or asparagine (condition similar to Fig. 3E), we observed a more than 2-fold decrease in S6K1 phosphorylation levels in cells infected with vΔVGF **(Fig. 4C).** We further confirmed these results as we observed an increase in mTORC1 S2448 phosphorylation in WT-VACV compared to vΔVGF infection **(Fig. 4D)**. These results confirm that in the absence of glutamine and asparagine, upon VACV infection, VGF is required to activate S6K1, the immediate downstream target of mTORC1 signaling. We further determined if the activation of mTORC1-S6K1 is necessary for the phosphorylation of CAD at S1859. On treatment of VACV infected HFFs with mTORC1 inhibitor rapamycin, at a condition that did not affect cell viability^53^, we observed a reduction in CAD S1859 phosphorylation in WT-VACV compared to vehicle or no treatment conditions **(Fig. 4E & 4F)**. These data indicate that in the absence of glutamine or asparagine, VGF is required to activate mTORC1-S6K1 for CAD phosphorylation, suggesting that the absence of these two amino acids in particular acts as a cue for VGF to regulate the mTORC1-CAD S1859 axis during VACV infection.

**Figure 4.**
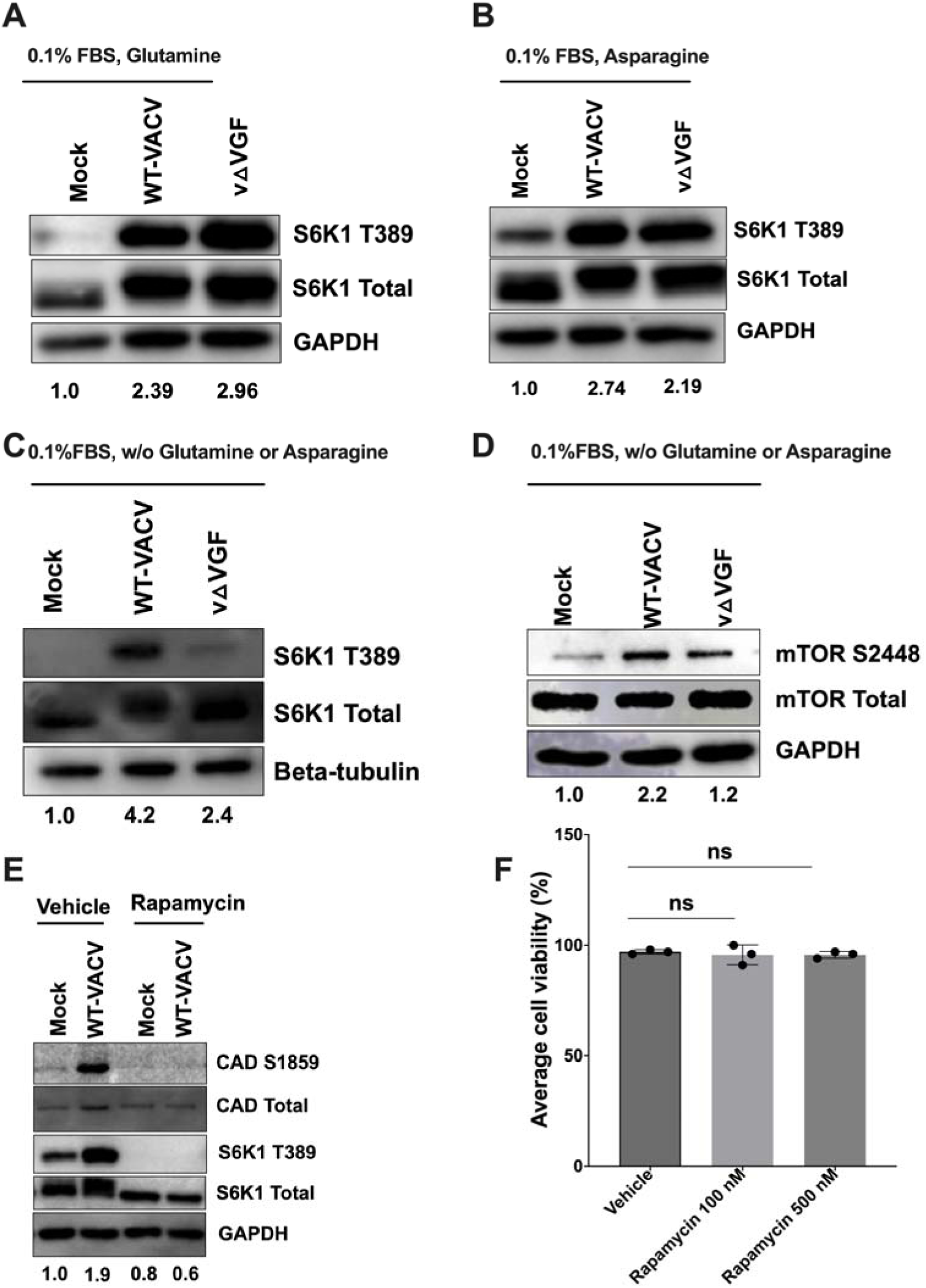
VGF is required to activate CAD through mTORC1-S6K1 in VACV infection. **(AB)** Role of VGF in activating mTORC1 downstream substrate S6K1 under different nutrient cues. HFFs were starved with 0.1% dialyzed FBS with either 2mM glutamine (**A**) or 2 mM asparagine **(B)** for 48 h prior to infection. Cells were infected with indicated viruses at an MOI of 5 for 8 h. **(C & D)** HFFs was starved in media with 0.1% dialyzed FBS and 2 mM asparagine, followed by replacement with media containing 0.1% dialyzed FBS lacking both glutamine and asparagine for approximately 16 h prior to infection. The infection media consisted of 0.1% dialyzed FBS without glutamine or asparagine. Infection was carried out with indicated viruses using MOI 5 for 8 h. **(E)** HFFs were infected in conditions described in Fig. 4CD. HFFs were either treated with vehicle (DMSO) or rapamycin (20 nM) 1 hpi and cell lysates were collected at 8 hpi. **(F)** HFFs were starved as described in Fig.4CD and treated with indicated concentration of rapamycin for 24 h. following starvation. Cell viability was measured using trypan blue staining. Representative images from two (D, E) or three (A, B, C,) biological replicates are shown. The numbers below the blot indicate the fold change of the indicated phosphorylated proteins based on the average intensities of Western blot analyses from two (D, E) or three (A, B, C) biological replicates, with each replicate normalized to the loading control of its respective blot, and then further normalized to its mock treatment. Western blot quantification was determined by NIH Image J.

### Expression and purification of recombinant VGF peptide

To investigate whether VGF in the absence of VACV infection could activate CAD and S6K1 signaling, we designed and expressed a recombinant VGF peptide using the processed, secreted form of VGF **(Fig. 5A)**. The VGF gene, designated as C11R, was codon-optimized, synthesized, and cloned into a plasmid vector pcDNA3.4, followed by expression using a mammalian expression system (CHO cells) to ensure proper post-translational modifications. A 6xhistidine tag (6H) was added to the peptide used for protein purification followed by size exclusion chromatography. The SEC-HPLC **(Fig. 5B)** and SDS-PAGE with Coomassie blue staining (**Fig. 5C**) were performed, which indicated that the peptide was highly pure at the expected molecular size (∼19 kDa). The identity of the recombinant peptide was further analyzed by sequence coverage after mass spectrometry analysis **(Fig. 5D)**.

**Figure 5.**
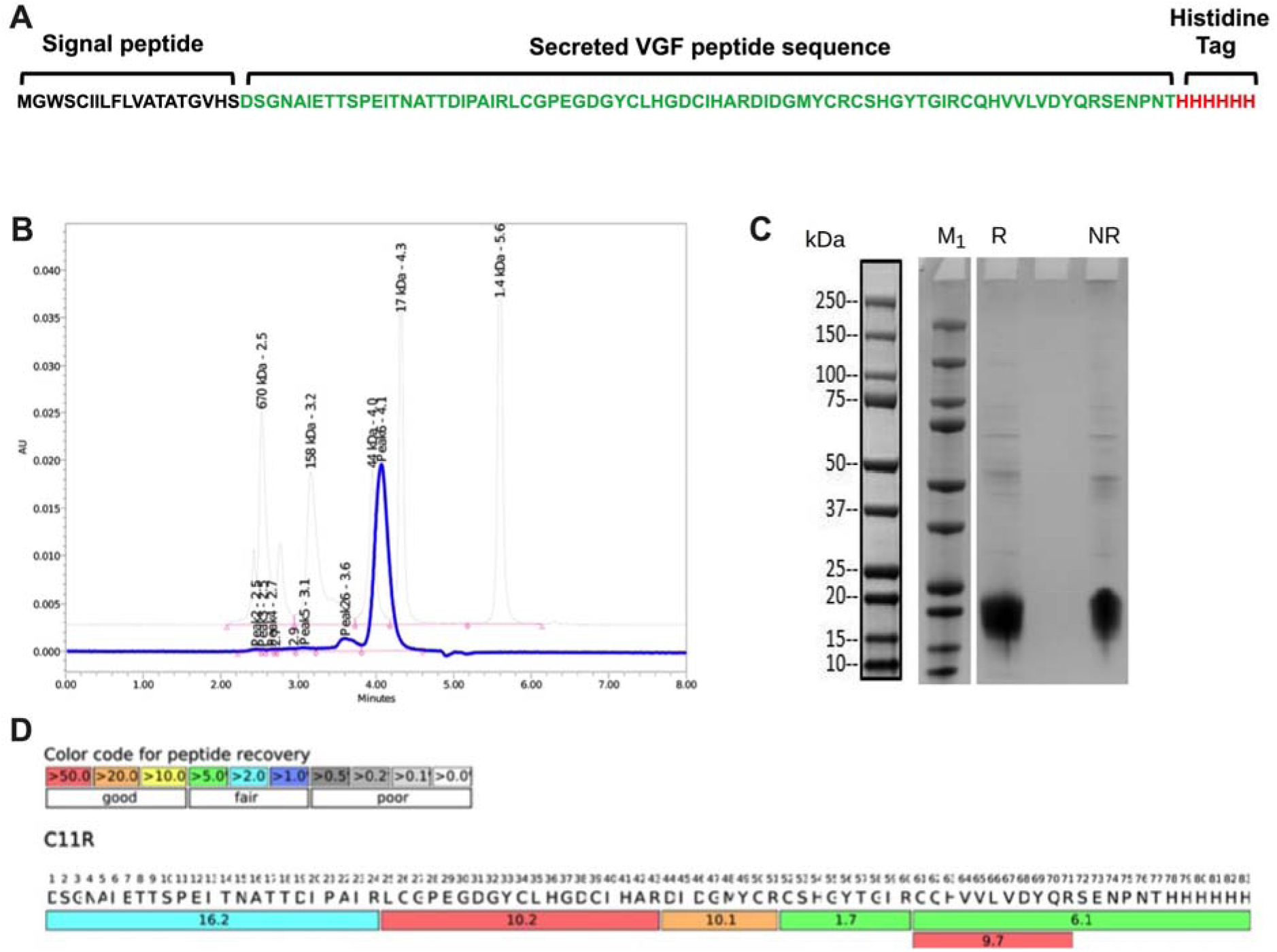
Expression, purification, and characterization of recombinant VGF peptide from a mammalian expression system. **(A)** Design of VGF sequence with the cleavable signal peptide at the N-terminal and a 6x Histidine tag at the C-terminal end. **(BC)** Purified VGF peptide was examined through **(B)** SEC-HPLC and **(C)** SDS-PAGE by Coomassie blue staining. M1: Protein marker, R- Reducing condition, NR-Non-reducing condition. **(D)** The sequence coverage was assessed with LC-MS after the purified VGF peptide was digested with trypsin.

### VGF peptide in the absence of VACV infection is insufficient to stimulate mTORC1-CAD S1859 in glutamine and asparagine deprived conditions

We next examined the activity of the recombinant VGF peptide by determining its ability to stimulate EGFR. We observed little stimulation of EGFR in mock and vΔVGF conditions compared to WT-VACV infection, confirming that VGF is needed to activate EGFR Y1068 **(Fig. 6A)**. The treatment of HFFs with VGF peptide alone could stimulate EGFR, albeit to a lesser extent than WT-VACV infection. However, treatment of HFFs with VGF peptide in the presence of vΔVGF infection, was able to fully rescue the effects on EGFR Y1068 phosphorylation, restoring it to levels similar to those observed with WT-VACV infection. On the contrary, while the EGF peptide alone could activate EGFR, its application in the presence of vΔVGF infection did not lead to full activation of the EGFR **(Fig. 6A)**. In fact, the infection condition reduced the phosphorylation levels to those similar to mock. The results demonstrated the biological activity of the recombinant VGF peptide. However, the data also indicates that, despite VGF being homologous to cellular EGF, viral infection creates a much more conducive environment for VGF activation of EGFR Y1068 than in the absence of infection, suggesting that EGF and VGF do not exhibit identical activities, possibly because of the acquirement of unique functions by VGF during poxvirus infection.

**Figure 6.**
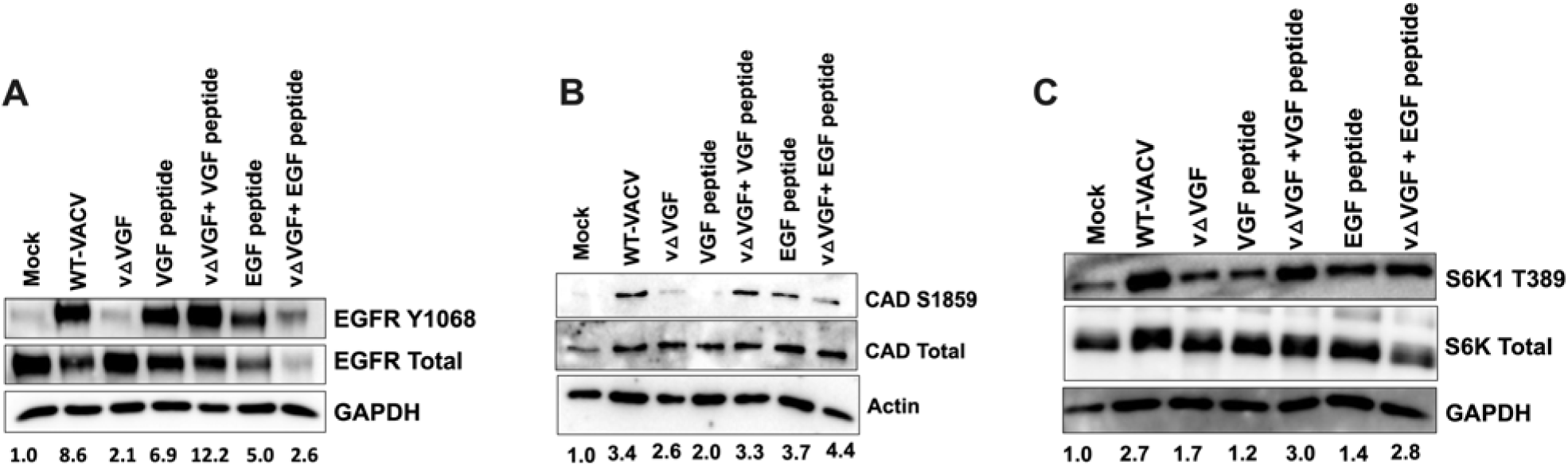
VGF peptide rescues vΔVGF reduction of CAD and S6K1 phosphorylation levels but not in the absence of infection. HFFs were starved with 0.1% dialyzed FBS without glutamine or asparagine as described in Fig.3E. Cells were infected with indicated viruses at an MOI of 5. VGF or EGF peptide was added at the time of infection for respective samples at a concentration of 5 µg/ml. Cells were treated for a total of 8 h prior to sample preparation for western blot analysis. EGFR **(A),** CAD S1859 **(B),** and S6K1 T389 **(C)** phosphorylation was detected by Western blotting analyses using indicated antibodies. Images presented are a representative from three biological replicates. The numbers below the blot indicate the average fold change of the phosphorylated protein of interest based of three biological replicates, with each replicate normalized to the loading control of its respective blot, and then further normalized to its mock treatment. Western blot quantification was determined by NIH Image J.

Subsequently, when determining the effects of the VGF peptide on the phosphorylation of S6K1 and CAD, we found that the VGF peptide alone was unable to increase phosphorylation levels of CAD S1859 and S6K1 T389 at a noticeable level **(Fig. 6BC)**. Phosphorylation levels for both proteins under VGF peptide treatment were similar or only very slightly higher than those in mock-treated cells. The EGF peptide, as well as the combinations of vΔVGF and VGF or vΔVGF and EGF were able to rescue the effects of vΔVGF on CAD and S6K1 phosphorylation levels. Overall, these data suggest that the VGF peptide, in the presence of vΔVGF infection, can compensate for the absence of VGF in activating CAD and mTORC1. However, VGF peptide alone, independent of infection, is unable to achieve the same effects. These discoveries suggest that other viral factors are needed to synergize with VGF for CAD and S6K1 activation upon VACV infection. It is interesting that total EGFR levels decreased in WT-VACV infection, or VGF/EGF treatment, which demands further investigations.

## Discussion

Our study demonstrates the importance of the *de novo* pyrimidine pathway during VACV replication and elucidates that the early VACV protein, VGF, is necessary for the stimulation of mTORC1-S6K1 and subsequent CAD phosphorylation at S1859 under nutrient-poor conditions **(Fig. 7)**. We highlight that in conditions replete with FBS, glutamine, or asparagine, HFFs exhibit similar levels of CAD phosphorylation (S1859 and T456, both of which are known to induce CAD activity), across uninfected and infected conditions. The absence of VGF does not significantly affect the phosphorylation of CAD, a crucial enzyme in the pyrimidine synthesis pathway in these conditions. However, VGF emerges as necessary for phosphorylating CAD at S1859, in conditions devoid of these two amino acids, and with minimal FBS **(Fig. 3)**.

**Figure 7.**
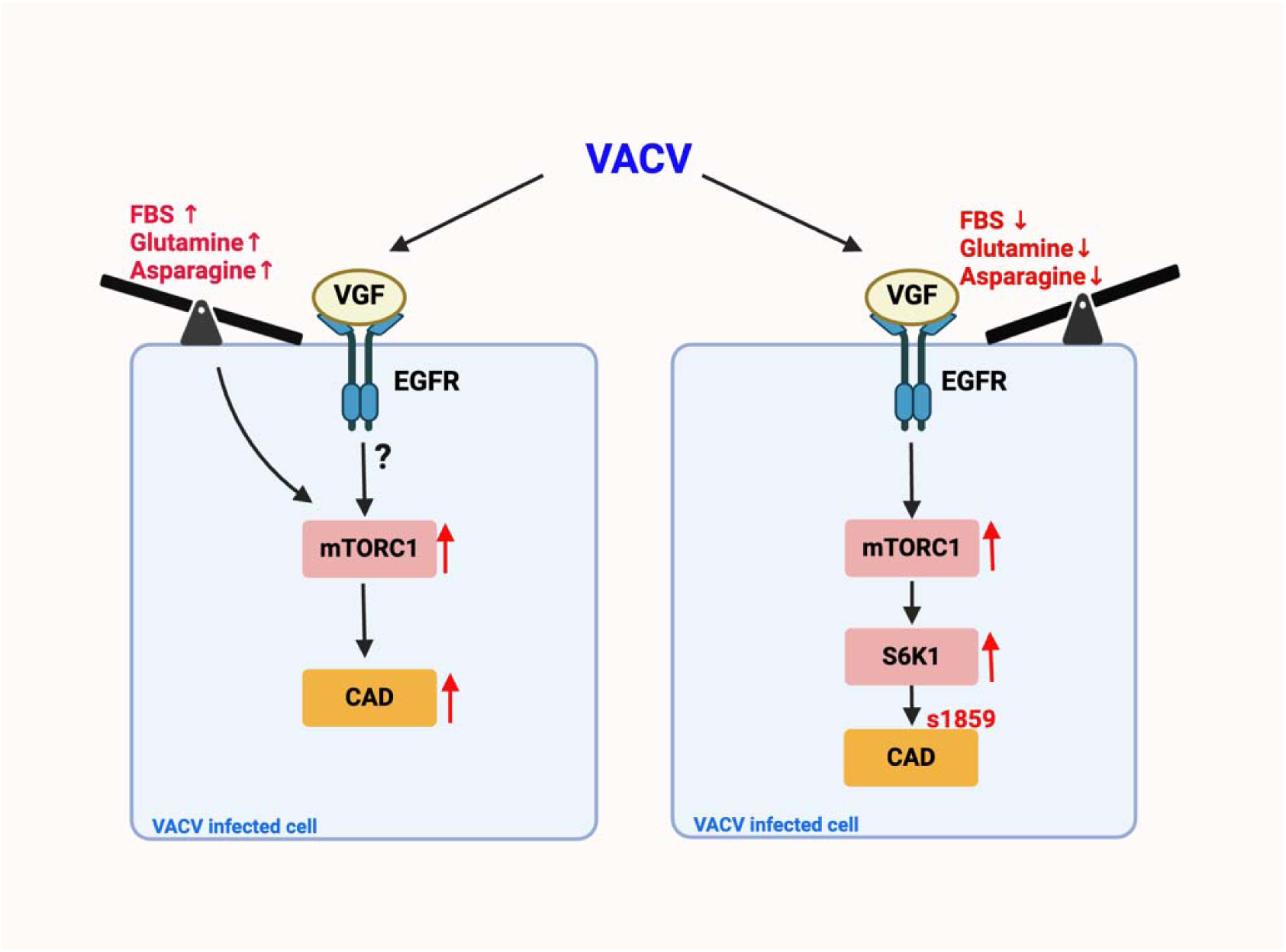
Proposed model of VACV’s regulation of the mTORC1-CAD axis under different nutrient cues. In nutrient-rich conditions, including glutamine, asparagine, and FBS, these nutrients can activate the CAD axis, thereby concealing the impact of VGF on the mTORC1-CAD axis. Conversely, in the absence of FBS, glutamine, and asparagine, VGF assumes a significant role in activating mTORC1 signaling to trigger CAD at serine1859, integral to the stimulation of the *de novo* pyrimidine pathway upon VACV infection.

The regulation of CAD is complex. CAD has multiple regulatory sites, with T456, S1859, and S1873 known to positively regulate CAD through the MAPK, mTORC1, or through PKC respectively^25^. We noted generally consistent levels of CAD phosphorylation at T456 across all nutrient conditions regardless of infection status **(Fig. 3)**. This highlights that T456 activation may be essential to maintain the fundamental cellular functions under various nutrient conditions irrespective of infection in HFFs. Though the VGF peptide alone could activate EGFR, it was insufficient to phosphorylate CAD and S6K1 independently of infection. However, the addition of the VGF peptide during vΔVGF infection offsets the reduction in CAD and S6K1 phosphorylation resulting from the absence of VGF in the vΔVGF infection condition **(Fig. 6)**. This highlights that VACV infection requires other viral factors, apart from VGF, to activate the mTORC1-CAD axis. Furthermore, compared to VGF’s ability to restore phosphorylation levels of EGFR upon vΔVGF infection, the decrease in EGFR phosphorylation observed with EGF peptide treatment during vΔVGF infection suggests that, despite sharing homology, VGF is more effective in stimulating EGFR activation during VACV infection. This may also partially explain the encoding of VGF in almost all poxviruses. Given EGFR’s pivotal role in initiating significant signaling cascades, these findings emphasize the necessity of investigating VGF specific mechanisms of activating the EGFR signaling cascade and subsequent effects during VACV infection^54^. These data also imply that VGF exhibits differential functionality compared to its cellular homolog. Understanding how VGF reprograms EGFR signaling would offer deeper insights into how poxviruses modify major host signaling pathways.

While we have gained a fresh perspective on how VACV modulates the CAD enzyme, further investigation is needed to elucidate other modes of CAD regulation. Apart from phosphorylation, other modes of regulation of proteins include acetylation, methylation, ubiquitinylation^55^. Whilst information of these post translational modifications (PTMs) of CAD remains scarce, there is evidence of other binding proteins like Rad9 and Rheb, that activate CAD activity^56, 57^. Both Rheb and Rad9 were shown to bind to CAD at the CPSII domain of CAD, away from the S1859 phosphorylation site, which is located between the DHOase and ATcase domain **(Fig. 3)**. The interaction of these proteins triggers the CPSIIase activity of CAD, while Rheb binding is also known to stimulate nucleotide pools. Additionally, there is evidence of Rheb being activated by EGFR signaling in certain tumors^58^. Given that VGF is a viral homolog of EGF, it is also plausible to hypothesize that VGF indirectly alters CAD through other CAD-interacting proteins like Rheb in conditions replete with nutrition. These reports, along with our findings, open up the possibility of VGF being involved in an alternate mode of regulating CAD, apart from phosphorylation at serine 1859, under conditions rich in glutamine and asparagine.

Accumulating evidence suggests that cancers consisting of KRAS mutations could be susceptible to the effects of DHODH enzyme inhibition^59^. Considering that KRAS activation is influenced by upstream growth factor signaling, there’s a possibility that VGF may play a role in regulating DHODH during VACV infection. Additional research is required to elucidate the precise mechanisms to identify if VACV regulates DHODH and UMPS, as well as to determine if VGF activates particular pathways to stimulate these enzymes under different nutritional conditions.

The *de novo* pyrimidine pathway not only supplies intermediates necessary for nucleic acid synthesis but also contributes to other cellular functions. UMP generated through this pathway can increase UDP-GlcNA levels, which are essential for O-GlcNAcylation of proteins. The O-GlcNAcylation of proteins serves as a nutritional sensor and stress receptor, responding to diverse environmental cues^60^. The importance of VGF-dependent CAD S1859 phosphorylation under nutrient-deficient conditions is suggestive of the potential involvement of CAD S1859 in activating the *de novo* pyrimidine pathway to increase UDP-GlcNA levels.

We also demonstrate that the phosphorylation of CAD S1859 is mediated by the phosphorylation of S6K1 at T389, a substrate of mTORC1 **(Fig. 4)** in conditions devoid of glutamine and asparagine upon VACV infection. mTORC1 serves as a central regulator of metabolism, integrating signals from diverse nutrients and growth factors to regulate both anabolic and catabolic processes^61^. Complete activation of mTORC1 cannot be achieved solely through growth factors; it needs both growth factor signaling and amino acid stimulation. Glutamine and asparagine have been shown to independently activate mTORC1 signaling^62^. The presence of these two amino acids in the infection media suggests that they might be sufficient to activate mTORC1 independently of VGF signaling upon VACV infection. Nevertheless, the decrease in S6K1 phosphorylation under conditions lacking asparagine and glutamine suggest that the signaling mediated by VGF to activate mTORC1 becomes critical during nutritional stress **(Fig. 4)**. Conditions with limited amino acids diminishes mTORC1 activity, leading to reduced cell growth and proliferation^63^. Previous studies have highlighted the continuous stimulation of mTORC1 activity by herpes simplex virus and human cytomegalovirus under amino acid deprivation conditions, providing initial insights into how viruses can regulate and maintain viral replication under physiological stress^64, 65^. However, whether poxviruses are able to activate mTORC1 during conditions of nutritional stress was unknown. Our research highlights that despite nutritional stress, such as the lack of glutamine, asparagine, and FBS, VACV retains the ability to activate the master regulator of metabolism, mTORC1, via its viral growth factor VGF. Additionally, the requirement for both amino acids and growth factors for the full activation of mTORC1 raises the question of whether VACV, together with VGF, utilizes another amino acid besides glutamine or asparagine for mTORC1 activation under conditions devoid of these two amino acids.

It has been reported that F17, synthesized late during VACV infection, dysregulates mTORC1^66^. While F17 is a virion core protein that is probably transferred to the host cell during entry, the virion containing F17 alone unlikely strongly activates mTORC1 within the first 8 hours post-infection in the absence of glutamine/asparagine. This is evident as we did not observe strong S6K1 activation in starved HFFs infected with vΔVGF. A possible explanation for this could be that VACV employs different viral proteins to activate distinct signaling pathway components in varying nutritional and physiological contexts. Alternatively, VGF and F17 may synergize to regulate mTORC1 signaling.

VACV has been utilized as an oncolytic agent for treating cancer cells, and there is increasing evidence of the benefits of prolonged starvation in enhancing the efficiency of oncolytic virotherapy in cancer treatment^37, 67^. Our discovery of the significance of VGF in stimulating mTORC1 under conditions of starvation provides insights into deciphering the molecular mechanisms by which VACV modulates metabolism in specific physiological contexts. This understanding can be useful in improving the effectiveness of VACV as an oncolytic virus against cancer cells. Additionally, tumor growth occurs in various nutritional environments, with different tumors experiencing distinct amino acid vulnerabilities^68, 69^. Understanding how VACV uses distinct signaling mechanisms in response to diverse nutritional cues will contribute to advancing the therapeutic potential of VACV as an oncolytic agent.

Unrestricted proliferation in conditions such as cancer makes nucleotide synthesis inhibition an appealing therapeutic strategy^70^. However, different viruses exhibit context-specific mechanisms for this uncontrolled proliferation^70^. Our study reveals new mechanisms of mTORC1 regulation via VGF early during infection. Additionally, there are multiple inhibitors of CAD that continue to face challenges due to off-target effects, underscoring the need for more potent and specific inhibitors of this enzyme^71^. Our aim to determine the specific regulatory sites of CAD essential for activation during VACV infection, aims to bridge gaps in our current understanding of this complex rate limiting enzyme for poxvirus specific therapy.

In summary, our study has revealed a mechanism utilized by VACV to activate mTORC1 under nutrient deficient conditions. We emphasize that viral early protein, VGF, is needed for activating CAD at S1859 in environments lacking glutamine and asparagine **(Fig. 7)**. These results suggest that the absence of glutamine and asparagine could possibly serve as a cue for activating VGF signaling upon VACV infection, consequently activating mTORC1 and ultimately leading to the activation of its downstream substrate CAD, which is required for the stimulation of the *de novo* pyrimidine pathway.

## Acknowledgements

We would like to express our gratitude to Dr. Bernard Moss and Dr. Nicholas Wallace for providing reagents, and to the members of the Yang Lab for their valuable comments. ChatGPT was used for assistance with grammar and writing style and we, the authors, take full responsibility of the content of this paper. ZY is supported, in part, by grants from the NIH (R01AI143709 and R01AI183580) and Texas A&M University Research Leadership Fellowship.

## References

1. Goodwin, C. M., Xu, S. & Munger, J. Stealing the Keys to the Kitchen: Viral Manipulation of the Host Cell Metabolic Network. Trends Microbiol. 23, 789–798 (2015).

2. Thaker, S. K., Ch’ng, J. & Christofk, H. R. Viral hijacking of cellular metabolism. BMC Biol. 17, 59 (2019).

3. Sanchez, E. L. & Lagunoff, M. Viral activation of cellular metabolism. Virology 479-480, 609–618 (2015).

4. Fontaine, K. A., Camarda, R. & Lagunoff, M. Vaccinia virus requires glutamine but not glucose for efficient replication. J. Virol. 88, 4366–4374 (2014).

5. Pant, A., Cao, S. & Yang, Z. Asparagine Is a Critical Limiting Metabolite for Vaccinia Virus Protein Synthesis during Glutamine Deprivation. J. Virol. 93, e01834–18 (2019).

6. Lagziel, S., Gottlieb, E. & Shlomi, T. Mind your media. Nat. Metab. 2, 1369–1372 (2020).

7. McFadden, G. Poxvirus tropism. Nat. Rev. Microbiol. 3, 201–213 (2005).

8. Lal, M. K., et al. Nutrient-Mediated Perception and Signalling in Human Metabolism: A Perspective of Nutrigenomics. Int. J. Mol. Sci. 23, 11305 (2022).

9. Future State of Smallpox Medical Countermeasures. (National Academies Press, Washington, D.C., 2024). doi:10.17226/27652.

10. Yang, Z., Monkeypox: A potential global threat? J Med Virol, 2022. 94(9): p. 4034–4036. - Google Search. https://www.google.com/search?q=Yang%2C+Z.%2C+Monkeypox%3A+A+potential+global+threat%3F+J+Med+Virol%2C+2022.+94(9)%3A+p.+4034-4036.&oq=Yang%2C+Z.%2C+Monkeypox%3A+A+potential+global+threat%3F+J+Med+Virol%2C+2022.+94(9)%3A+p.+4034-4036.&gs_lcrp=EgZjaHJvbWUyBggAEEUYOdIBBzQ4MWowajeoAgCwAgA&sourceid=chrome&ie=UTF-8.

11. Durski, K. N. Emergence of Monkeypox — West and Central Africa, 1970–2017. MMWR Morb. Mortal. Wkly. Rep. 67, (2018).

12. Yang, Z., Gray, M. & Winter, L. Why do poxviruses still matter? Cell Biosci. 11, 96 (2021).

13. Verardi, P. H., Titong, A. & Hagen, C. J. A vaccinia virus renaissance. Hum. Vaccines Immunother. 8, 961–970 (2012).

14. Hendrickson, R. C., Wang, C., Hatcher, E. L. & Lefkowitz, E. J. Orthopoxvirus Genome Evolution: The Role of Gene Loss. Viruses 2, 1933–1967 (2010).

15. Chang, W., Lim, J. G., Hellström, I. & Gentry, L. E. Characterization of vaccinia virus growth factor biosynthetic pathway with an antipeptide antiserum. J. Virol. 62, 1080–1083 (1988).

16. Blomquist, M. C., Hunt, L. T. & Barker, W. C. Vaccinia virus 19-kilodalton protein: relationship to several mammalian proteins, including two growth factors. Proc. Natl. Acad. Sci. U. S. A. 81, 7363–7367 (1984).

17. Twardzik, D. R., Brown, J. P., Ranchalis, J. E., Todaro, G. J. & Moss, B. Vaccinia virus-infected cells release a novel polypeptide functionally related to transforming and epidermal growth factors. Proc. Natl. Acad. Sci. U. S. A. 82, 5300–5304 (1985).

18. Yang, Z., Bruno, D. P., Martens, C. A., Porcella, S. F. & Moss, B. Simultaneous high-resolution analysis of vaccinia virus and host cell transcriptomes by deep RNA sequencing. Proc. Natl. Acad. Sci. U. S. A. 107, 11513–11518 (2010).

19. Beerli, C. et al. Vaccinia virus hijacks EGFR signalling to enhance virus spread through rapid and directed infected cell motility. Nat. Microbiol. 4, 216–225 (2019).

20. Mazzon, M., Castro, C., Roberts, L. D., Griffin, J. L. & Smith, G. L. A role for vaccinia virus protein C16 in reprogramming cellular energy metabolism. J. Gen. Virol. 96, 395–407 (2015).

21. Pant, A., Dsouza, L., Cao, S., Peng, C. & Yang, Z. Viral growth factor- and STAT3 signaling-dependent elevation of the TCA cycle intermediate levels during vaccinia virus infection. PLOS Pathog. 17, e1009303 (2021).

22. De novo Fatty Acid Biosynthesis Contributes Significantly to Establishment of a Bioenergetically Favorable Environment for Vaccinia Virus Infection | PLOS Pathogens. https://journals.plos.org/plospathogens/article?id=10.1371/journal.ppat.1004021.

23. Pant, A., Brahim Belhaouari, D., Dsouza, L. & Yang, Z. Upregulation of ATP Citrate Lyase Phosphorylation and Neutral Lipid Synthesis through Viral Growth Factor Signaling during Vaccinia Virus Infection. Preprint at 10.1101/2023.09.21.558916 (2023).

24. Huang, M. & Graves, L. M. De novo synthesis of pyrimidine nucleotides; emerging interfaces with signal transduction pathways. Cell. Mol. Life Sci. CMLS 60, 321–336 (2003).

25. del Caño-Ochoa, F., Moreno-Morcillo, M. & Ramón-Maiques, S. CAD, A Multienzymatic Protein at the Head of de Novo Pyrimidine Biosynthesis. in Macromolecular Protein Complexes II: Structure and Function (eds. Harris, J. R. & Marles-Wright, J.) vol. 93 505–538 (Springer International Publishing, Cham, 2019).

26. Walter, M. & Herr, P. Re-Discovery of Pyrimidine Salvage as Target in Cancer Therapy. Cells 11, 739 (2022).

27. Lane, A. N. & Fan, T. W.-M. Regulation of mammalian nucleotide metabolism and biosynthesis. Nucleic Acids Res. 43, 2466–2485 (2015).

28. Cytomegalovirus-mediated activation of pyrimidine biosynthesis drives UDP–sugar synthesis to support viral protein glycosylation | PNAS. https://www.pnas.org/doi/full/10.1073/pnas.1415864111.

29. Tang, N. et al. Newcastle Disease Virus Manipulates Mitochondrial MTHFD2-Mediated Nucleotide Metabolism for Virus Replication. J. Virol. 97, e0001623 (2023).

30. Valle-Casuso, J.-C. et al. Replication of Equine arteritis virus is efficiently suppressed by purine and pyrimidine biosynthesis inhibitors. Sci. Rep. 10, 10100 (2020).

31. Katsafanas, G. C., Grem, J. L., Blough, H. A. & Moss, B. Inhibition of vaccinia virus replication by N-(phosphonoacetyl)-L-aspartate: differential effects on viral gene expression result from a reduced pyrimidine nucleotide pool. Virology 236, 177–187 (1997).

32. Evans, D. R. & Guy, H. I. Mammalian Pyrimidine Biosynthesis: Fresh Insights into an Ancient Pathway *. J. Biol. Chem. 279, 33035–33038 (2004).

33. Sigoillot, F. D. et al. Protein kinase C modulates the up-regulation of the pyrimidine biosynthetic complex, CAD, by MAP kinase. Front. Biosci.-Landmark 12, 3892–3898 (2007).

34. Sigoillot, F. D. et al. Nuclear Localization and Mitogen-activated Protein Kinase Phosphorylationof the Multifunctional ProteinCAD. J. Biol. Chem. 280, 25611–25620 (2005).

35. Szwed, A., Kim, E. & Jacinto, E. Regulation and metabolic functions of mTORC1 and mTORC2. Physiol. Rev. 101, 1371–1426 (2021).

36. Saxton, R. A. & Sabatini, D. M. mTOR Signaling in Growth, Metabolism, and Disease. Cell 168, 960–976 (2017).

37. Li, M., Zhang, M., Ye, Q., Liu, Y. & Qian, W. Preclinical and clinical trials of oncolytic vaccinia virus in cancer immunotherapy: a comprehensive review. Cancer Biol. Med. 20, 646–661 (2023).

38. Yaghchi, C. A., Zhang, Z., Alusi, G., Lemoine, N. R. & Wang, Y. Vaccinia virus, a promising new therapeutic agent for pancreatic cancer. Immunotherapy 7, 1249–1258 (2015).

39. Cotter, C. A., Earl, P. L., Wyatt, L. S. & Moss, B. Preparation of Cell Cultures and Vaccinia Virus Stocks. Curr. Protoc. Protein Sci. 89, 5.12.1–5.12.18 (2017).

40. Baer, A. & Kehn-Hall, K. Viral Concentration Determination Through Plaque Assays: Using Traditional and Novel Overlay Systems. J. Vis. Exp. JoVE 52065 (2014) doi:10.3791/52065.

41. Strober, W. Trypan Blue Exclusion Test of Cell Viability. Curr. Protoc. Immunol. 111, A3.B.1–A3.B.3 (2015).

42. Li, G., Li, D., Wang, T. & He, S. Pyrimidine Biosynthetic Enzyme CAD: Its Function, Regulation, and Diagnostic Potential. Int. J. Mol. Sci. 22, 10253 (2021).

43. Collins, K. D. & Stark, G. R. Aspartate Transcarbamylase. J. Biol. Chem. 246, 6599–6605 (1971).

44. Teschner, S. & Burst, V. Leflunomide: A Drug with a Potential Beyond Rheumatology. Immunotherapy 2, 637–650 (2010).

45. Ringer, D. P., Howell, B. A. & Etheredge, J. L. Alteration in de novo pyrimidine biosynthesis during uridine reversal of pyrazofurin-inhibited dna synthesis. J. Biochem. Toxicol. 6, 19–27 (1991).

46. Cadman, E. C., Dix, D. E. & Handschumacher, R. E. Clinical, biological, and biochemical effect of pyrazofurin. Cancer Res. 38, 682–688 (1978).

47. Greene, S., Watanabe, K., Braatz-Trulson, J. & Lou, L. Inhibition of dihydroorotate dehydrogenase by the immunosuppressive agent leflunomide. Biochem. Pharmacol. 50, 861–867 (1995).

48. Greseth, M. D. & Traktman, P. The Life Cycle of the Vaccinia Virus Genome. Annu. Rev. Virol. 9, 239–259 (2022).

49. Villa, E., Ali, E., Sahu, U. & Ben-Sahra, I. Cancer Cells Tune the Signaling Pathways to Empower de Novo Synthesis of Nucleotides. Cancers 11, 688 (2019).

50. Robitaille, A. M. et al. Quantitative Phosphoproteomics Reveal mTORC1 Activates de Novo Pyrimidine Synthesis. Science 339, 1320–1323 (2013).

51. Ben-Sahra, I., Howell, J. J., Asara, J. M. & Manning, B. D. Stimulation of de Novo Pyrimidine Synthesis by Growth Signaling Through mTOR and S6K1. Science 339, 1323–1328 (2013).

52. Battaglioni, S., Benjamin, D., Wälchli, M., Maier, T. & Hall, M. N. mTOR substrate phosphorylation in growth control. Cell 185, 1814–1836 (2022).

53. Lamming, D. W. Inhibition of the Mechanistic Target of Rapamycin (mTOR)–Rapamycin and Beyond. Cold Spring Harb. Perspect. Med. 6, a025924 (2016).

54. Oda, K., Matsuoka, Y., Funahashi, A. & Kitano, H. A comprehensive pathway map of epidermal growth factor receptor signaling. Mol. Syst. Biol. 1, 2005.0010 (2005).

55. Lee, M. J. & Yaffe, M. B. Protein Regulation in Signal Transduction. Cold Spring Harb. Perspect. Biol. 8, a005918 (2016).

56. Lindsey-Boltz, L. A., Wauson, E. M., Graves, L. M. & Sancar, A. The human Rad9 checkpoint protein stimulates the carbamoyl phosphate synthetase activity of the multifunctional protein CAD. Nucleic Acids Res. 32, 4524–4530 (2004).

57. Sato, T. et al. Rheb Protein Binds CAD (Carbamoyl-phosphate Synthetase 2, Aspartate Transcarbamoylase, and Dihydroorotase) Protein in a GTP- and Effector Domain-dependent Manner and Influences Its Cellular Localization and Carbamoyl-phosphate Synthetase (CPSase) Activity. J. Biol. Chem. 290, 1096–1105 (2015).

58. Uribe, M. L., Marrocco, I. & Yarden, Y. EGFR in Cancer: Signaling Mechanisms, Drugs, and Acquired Resistance. Cancers 13, 2748 (2021).

59. Koundinya, M. et al. Dependence on the Pyrimidine Biosynthetic Enzyme DHODH Is a Synthetic Lethal Vulnerability in Mutant KRAS-Driven Cancers. Cell Chem. Biol. 25, 705–717.e11 (2018).

60. Liu, Y. et al. O-GlcNAcylation: the “stress and nutrition receptor” in cell stress response. Cell Stress Chaperones 26, 297–309 (2021).

61. Goul, C., Peruzzo, R. & Zoncu, R. The molecular basis of nutrient sensing and signalling by mTORC1 in metabolism regulation and disease. Nat. Rev. Mol. Cell Biol. 24, 857–875 (2023).

62. Meng, D. et al. Glutamine and asparagine activate mTORC1 independently of Rag GTPases. J. Biol. Chem. 295, 2890–2899 (2020).

63. Liu, G. Y. & Sabatini, D. M. mTOR at the nexus of nutrition, growth, ageing and disease. Nat. Rev. Mol. Cell Biol. 21, 183–203 (2020).

64. Clippinger, A. J., Maguire, T. G. & Alwine, J. C. Human Cytomegalovirus Infection Maintains mTOR Activity and Its Perinuclear Localization during Amino Acid Deprivation ▿. J. Virol. 85, 9369–9376 (2011).

65. Vink, E. I., Lee, S., Smiley, J. R. & Mohr, I. Remodeling mTORC1 Responsiveness to Amino Acids by the Herpes Simplex Virus UL46 and Us3 Gene Products Supports Replication during Nutrient Insufficiency. J. Virol. 92, e01377–18 (2018).

66. Meade, N., King, M., Munger, J. & Walsh, D. mTOR Dysregulation by Vaccinia Virus F17 Controls Multiple Processes with Varying Roles in Infection. J. Virol. 93, e00784–19 (2019).

67. Scheubeck, G. et al. Starvation-Induced Differential Virotherapy Using an Oncolytic Measles Vaccine Virus. Viruses 11, 614 (2019).

68. Lyssiotis, C. A. & Kimmelman, A. C. Metabolic Interactions in the Tumor Microenvironment. Trends Cell Biol. 27, 863–875 (2017).

69. Pathria, G. & Ronai, Z. A. Harnessing the Co-vulnerabilities of Amino Acid-Restricted Cancers. Cell Metab. 33, 9–20 (2021).

70. Ariav, Y., Ch’ng, J. H., Christofk, H. R., Ron-Harel, N. & Erez, A. Targeting nucleotide metabolism as the nexus of viral infections, cancer, and the immune response. Sci. Adv. 7, eabg6165 (2021).

71. Wang, W., Cui, J., Ma, H., Lu, W. & Huang, J. Targeting Pyrimidine Metabolism in the Era of Precision Cancer Medicine. Front. Oncol. 11, 684961 (2021).

